# LKB1 negatively regulates AKT1 signaling via DBC1 and TRB3

**DOI:** 10.1101/691402

**Authors:** Zarka Sarwar, Sameer Bhat, Qaaifah Gillani, Irfana Reshi, Misbah Un Nisa, Guillaume Adelmant, Jarrod Marto, Shaida Andrabi

## Abstract

DBC1 plays a critical role in various cellular functions notably cell proliferation, transcription, histone modification and adipogenesis. Current reports about the role of DBC1 in tumorigenesis are paradoxical and designate DBC1 both as a tumor suppressor or an oncogene. Here, using small T antigen of polyoma virus (PyST) as a tool, we have delineated a signaling mechanism that connects LKB1 to AKT1 via DBC1. We report that PyST associates with DBC1 and leads to its down-regulation. Our results also show that PyST expression promotes LKB1 activation which in turn leads to in the downregulation of DBC1 protein. Absence of DBC1 results in transcriptional upregulation and consequently enhanced protein levels of TRB3. TRB3 sequesters AKT1, and consequently the phosphorylation and activity of AKT1 is compromised. This ultimately results in inactivation of pro-survival pathways triggered via AKT1 signaling. Our studies thus provide an insight into a signaling pathway that connects LKB1, DBC1, TRB3 and AKT1.

## Introduction

Deleted in Breast Cancer 1 (DBC1), also known as CCAR2, was first identified in 2002 and was found to be deleted in human breast carcinomas (1). Several studies later reported that DBC1 mRNA is upregulated in breast carcinomas (2). Since then, the reported functions of DBC1 as a potential oncogene or tumor suppressor have been challenged by observation of its respective up and down-regulation in various cancers (3). For these reasons, the role of DBC1 in tumorigenesis remains undefined.

LKB1 (also called STK11), is a ubiquitously expressed serine/threonine kinase and is known to phosphorylate the family of 14 kinases referred to as AMPK-related kinases (ARKs) and is therefore termed a “master kinase” (4). LKB1 is implicated in diverse cellular functions particularly energy metabolism, cell cycle arrest, cell polarity and tumorigenesis. Germline mutations in human LKB1 are the main cause of Peutz-Jeghers Syndrome (PJS) (5) and such patients are at very high risk of developing hemartomas that can develop into tumors (6). While the regulation of AMPK activity by LKB1 is very well studied, less is known about other kinases and signaling pathways regulated by LKB1. In this paper, we show that DBC1 is a novel target of LKB1 that does not involve AMPK.

Polyoma viruses are a group of small DNA-based viruses and are potentially oncogenic in a broad variety of hosts. Expression of the small T antigen of the early region of polyoma virus in mammalian cells is known to induce apoptosis. This property of PyST is dependent upon its ability to bind the cellular serine/threonine phosphatase, PP2A (7). PyST is a small protein of 195 amino acids (M. wt. 20 kDa) and consists of an N-terminal J domain and a C-terminal domain. The N-terminal domain consists of an HPDKGG motif which is common to all T antigens of the polyoma virus family and binds heat shock proteins (HSPs). The C-terminal domain has two small cysteine motifs that have structural roles and also help bind PP2A (Fig. S1c). In subsequent studies, using inducible PyST expressing U2OS cell lines, it has been shown that expression of PyST causes a striking phenotype in the host cells. These cells get arrested in mitosis as can be seen by extensive rounding up, which is a hallmark of cells undergoing mitosis (8). Prolonged mitotic arrest is followed by overwhelming apoptosis within 3-4 days. Cells arrested in mitosis show various abnormalities, like multipolar spindles, lack of chromosomal congression and improper attachment of the chromosomes on the spindle (8). These cells also have activated “Spindle Assembly Checkpoint” (SAC) as is evident by the activation of various markers like Bub1 and BubR1 (8). However, the detailed mechanism that leads to this PyST induced phenotype and mitotic arrest is not yet known.

In pursuit of understanding these complexities, we wanted to identify the cellular proteins that interact with PyST. Using PyST expressing stable cell line, we show that DBC1 is an important component of cell cycle regulation that promotes mitotic exit under conditions of cellular stress as well as cellular transformation. More importantly, we also report that LKB1 negatively regulates AKT1 activity through DBC1 and Tribbles3 (TRB3). We therefore propose that down-regulation of DBC1 may be one of the mechanisms by which LKB1 exerts its tumor suppressor functions.

## Results

### Polyoma small T antigen interacts with DBC1 and causes its down-regulation

To understand the detailed mechanism of PyST induced mitotic arrest and apoptosis, we wanted to find out the potential cellular interacting partners for small T antigen. For that purpose, we made use of inducible stable U2OS cell lines expressing PyST having HA and Flag tags at the C-terminus. These cell lines, designated as pTREX-PyST-HA-Flag have been previously described (8). Addition of doxycycline causes expression of PyST in these cells and leads to remarkable rounding up phenotype, a feature usually associated with cells undergoing mitosis. To find out PyST interacting proteins, we performed tandem affinity purification (TAP) on the cell lysates obtained at about 20 hours of doxycycline addition, using anti-HA and anti-Flag antibodies respectively. A small aliquot of the immunoprecipitates was run on SDS-PAGE, and analyzed by silver staining. In this gel, bands for PyST and some of its well-known binding partners like A and C subunits of protein phosphatase 2A (PP2A) and HSP70 were clearly seen (Fig. 1a). The rest of the immunoprecipitated mixture was subjected to LC-MS-MS analysis. DBC1 (CCAR2) was found to be one of the many interacting partners of PyST and showed significant coverage of peptides in these mass spectrometry results (Fig. 1b and S1 a, b). To confirm that PyST and DBC1 directly interact with each other in the cells, we co-expressed PyST-HA-Flag and myc-DBC in 293T cells and subjected the lysates to immunoprecipitation by anti Flag (for PyST) antibodies. Western blotting results using anti myc antibody for DBC1 confirmed that DBC1 interacts with PyST (Fig.1c). Further support for the interaction of PyST and DBC1 came from immunofluorescence microscopy results. DBC1 was exclusively nuclear in its localization, as is already known. In presence of doxycycline (+dox), PyST was expressed and found predominantly in the nucleus, co-localizing with the DBC1 (Fig. 1d). To study the impact of PyST interaction on DBC1 expression and functions, we transiently transfected both these proteins in 293T cells. It was observed that PyST co-expression causes down-regulation of both endogenous as well as exogenously expressing protein levels of DBC1 (Fig. 1e and 1f). To find out the particular domain of PyST that is required for down-regulation of DBC1, we co-transfected myc-DBC1 with wild type PyST, PyST-BC1075 (PP2A mutant) and PyST-D44N (J domain mutant) in 293T cells. PyST-D44N contains a D^44^→N point mutation in the HPDKGG motif and thus cannot bind heat shock proteins (HSPs). Western blotting results showed that the down-regulation of DBC1 protein levels occurred when it was co-transfected with wild type and PyST-BC1075 but not with PyST-D44N (Fig. 1g). Further support for the PP2A independent role of PyST in causing DBC1 down-regulation came from the results showing that PP2A inhibitor okadaic acid also did not affect the protein levels of DBC1 in 293T cells (Fig. S1d). These results thus suggested the role of HSPs in regulating DBC1 protein levels. This is not entirely surprising given that the J domain of PyST is known to interact with YAP1 and TAZ, proteins having known oncogenic role (9, 10).

**Fig. 1:**
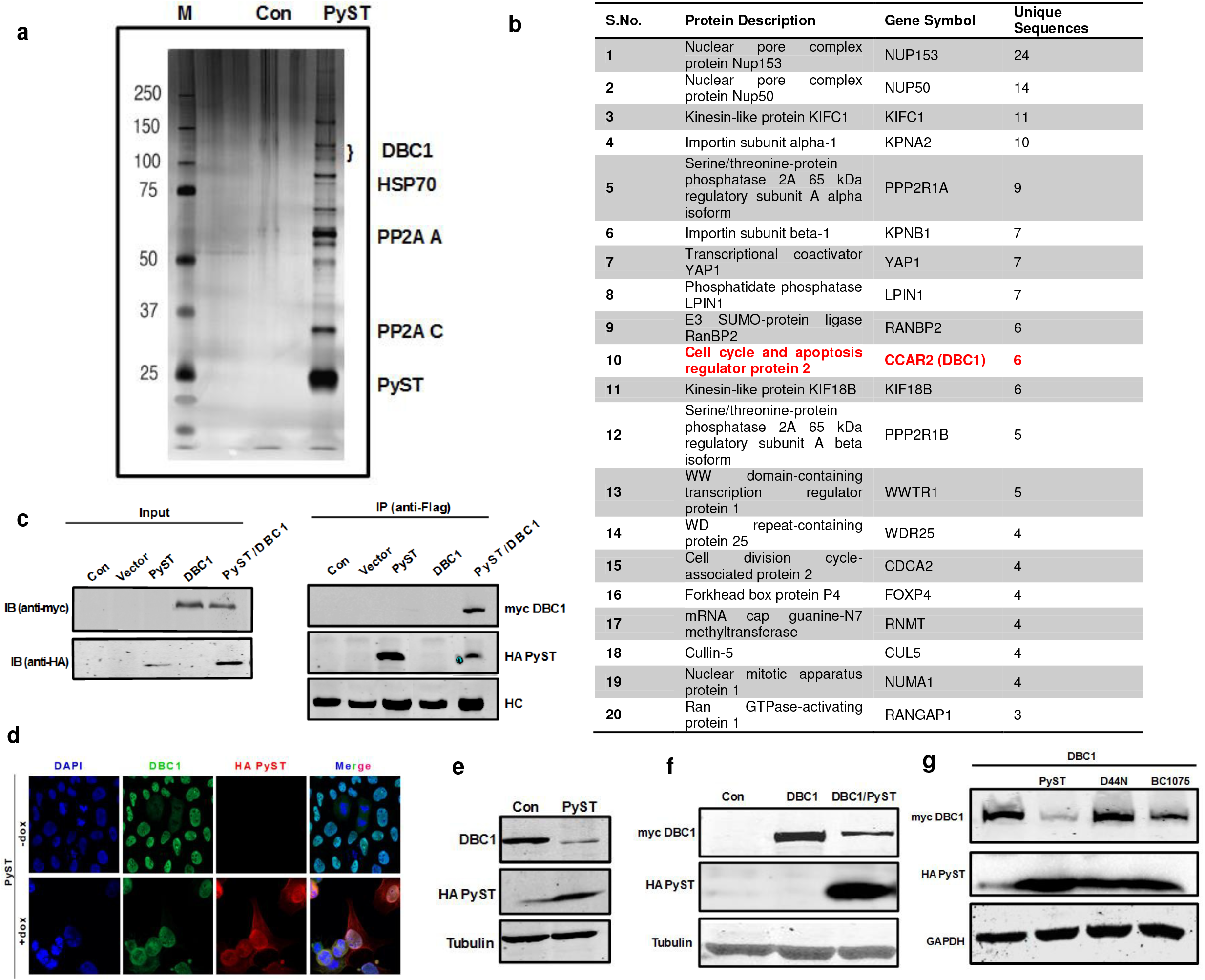
Polyoma small T (PyST) interacts with DBC1 and down-regulates its protein levels. (a) PyST associated complexes isolated from pTREX-PyST-HA-FLAG cell lines by tandem affinity purification (TAP) using consecutive anti-FLAG and anti-HA affinity matrices, were run on SDS-PAGE and visualized by silver staining. (b) Table showing proteins identified by 3 or more gene-unique peptides following LC-MS-MS analysis of PyST complex. (c) 293T cells were co-transfected with pcDNA3-PyST-HA-Flag, myc-DBC1 and empty vector (pcDNA3). PyST-HA-Flag was immunoprecipitated with anti-Flag antibody and subjected to SDS-PAGE and immunoblotting (IB) with indicated antibodies. HC: heavy chain of anti-Flag antibody used for immunoprecipitation. (d) Co-localization results of PyST (red) and DBC1 (green) using immunofluorescence microscopy obtained at about 24hrs of doxycycline addition. Nuclei were stained with DAPI. (e) 293T cells were transfected with pCDNA3-PyST-HA-Flag. After 48h, cell extracts were blotted for endogenous DBC1 using anti-DBC1 antibody and anti-HA antibody for PyST. (f) 293T cells were co-transfected with PyST-HA-Flag and myc-DBC1 constructs. Cell lysates were blotted for exogenously expressed DBC1 using anti-myc tag antibody and PyST using anti-HA antibody. (g) Degradation of DBC1 is dependent upon J domain of PyST. 293T cells were co-transfected with myc-DBC1 and different PyST-HA-Flag constructs like wild type PyST, BC1075 (PP2A mutant) or D44N (J domain mutant) constructs. Western blot analysis for DBC1 and PyST was carried out using indicated antibodies.

### Polyoma small T antigen causes post-translational down-regulation of DBC1

We wanted to find out whether the down-regulation of DBC1 was also valid in inducible PyST expressing U2OS cell lines (pTREX-PyST-HA-Flag) as mentioned above. Upon doxycycline addition, we saw time dependent decrease in the endogenous protein levels of DBC1 (Fig. 2a). To find out the impact of PyST on exogenous DBC1, we made double stable cell line expressing both PyST and DBC1 (pTREX-PyST-DBC1). Expression of PyST and DBC1 was confirmed by Western blotting (Fig. S2a). Similar results were obtained in these cell lines as well (Fig. 2b). Using RT-qPCR we observed that PyST expression had almost no effect on the mRNA levels of DBC1, implicating that down-regulation of DBC1 occurs at translational or post-translational level but not at transcriptional level (Fig. 2c).

**Fig. 2:**
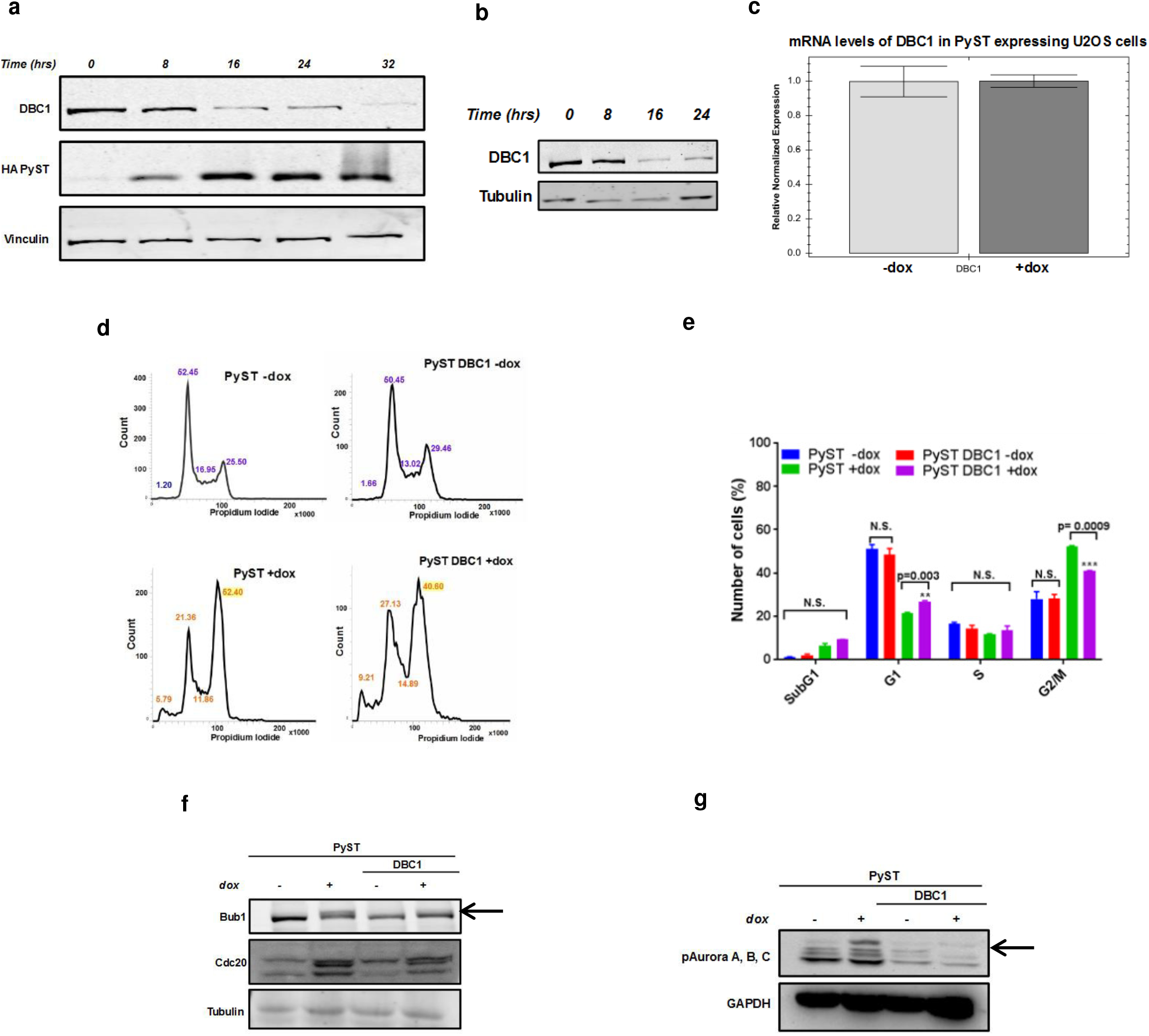
Small T causes post-translational down regulation of endogenous DBC1 protein levels and affects mitotic progression of cells. (a) Western blot analysis of PyST-HA-Flag expressing cell extracts at different time points of doxycycline addition using anti-DBC1 and anti-HA antibodies for detecting DBC1 (endogenous) and PyST respectively. (b) Effect of PyST overexpression on stably expressing exogenous myc-DBC1 protein levels as seen by Western blotting. (c) Effects of overexpression of PyST on DBC1 mRNA at about 24 hours of doxycycline addition. Following cDNA synthesis, RT-qPCR was carried out using primers for DBC1 and Actin. (d) FACS analysis of cells expressing PyST and DBC1 at 24 hours of doxycycline addition. (e) Statistical significance of FACS data was calculated using GraphPad Prism 7.0 (*P<0.05, **P<0.01, ***P<0.001). (f, g) DBC1 co-expression reduced protein levels of Bub1, Cdc20 and pAurora A, B, C in PyST expressing cells, following doxycycline addition for about 24 hours. PyST expression caused band shift in Bub1 and Aurora Kinase indicating phosphorylation.

### DBC1 plays an important role in mitotic exit

Since PyST overwhelmingly induces mitotic arrest and apoptosis in host cells and it also interacts with DBC1, we investigated the impact of DBC1 over-expression on PyST induced mitotic arrest in U2OS cells. We used pTREX-PyST-DBC1 (double stable) cell lines for cell cycle analysis using flow cytometry. Consistent with published reports (8), expression of PyST in U2OS cell lines induced a very robust mitotic arrest, (Fig. 2d) at 24 hours of PyST induction, as indicated by increased G2/M peak. On the other hand, PyST cells overexpressing DBC1, exhibited a decreased G2/M peak vis-à-vis an increase in G0/G1 peak (Fig. 2d, e). These results suggest that DBC1 somehow helps the cells overcome PyST induced mitotic arrest. This was also supported by Western blotting analysis of Bub1 kinase, Cdc20 and pAurora A, B, C in these cell lines. In pTREX-PyST-DBC1 cells (induced for PyST expression), protein levels of Cdc20 and phosphorylation levels of Bub1 and Aurora A, B, C levels were reduced (Fig. 2f and 2g). These results indicated that DBC1 expression aids in mitotic exit of PyST expressing cells.

Next, we wanted to test whether over-expression of DBC1 in PyST expressing cells (pTREX-PyST-DBC1) will have any effect on PyST mediated apoptosis following mitotic arrest. Crystal violet staining results on day 6 showed that DBC1 over-expression promoted the survival and growth of cells in presence of PyST expression (Fig. S2b). Similar down-regulation of DBC1 was seen when U2OS cells were treated with nocodazole and paclitaxel, drugs that arrest cells in mitosis (Fig. S2c). This implies a potential connection between mitotic arrest and decrease in DBC1 protein levels.

### PyST induces energy deficit and activates LKB1-AMPK pathway

The signaling cascades that lead to PyST induced mitotic catastrophe remain poorly understood. The pathways that are known to be affected the PyST include inhibition of PP2A, activation of p38 and ADP ribosylation of various cellular proteins (7). Since ADP Ribosylation induces extreme energy deficit, it was expected to activate the energy sensing machinery in the cells. LKB1 and AMPK are the major cellular energy sensors. We therefore wanted to check the status of LKB1 and AMPK activation in PyST cell lines. Using phospho S428 LKB1 antibody, we observed that these cells had elevated levels of phosphorylated LKB1 (Fig. 3a). Further confirmation came from the activation of AMP kinase, which is phosphorylated by LKB1 at T172 (Fig. 3b). To investigate direct involvement of LKB1/AMPK in regulation of DBC1, we analyzed the protein sequence of DBC1 using Scansite (www.scansite3.mit.edu) Results revealed a strong substrate consensus site for possible AMP Kinase phosphorylation (at T897) (Fig. S3 a). Since LKB1 is a well-known upstream kinase of AMPK, we wanted to test whether both LKB1 and AMPK would have any impact on DBC1 protein levels. Results showed that AMPK was not affecting DBC1 while LKB1 caused down-regulation of endogenous as well as exogenous (myc-DBC1) DBC1 (Fig. 3c, 3d respectively). To find out whether the T897 residue of DBC1 (predicted phosphorylation site for AMPK) has any importance in the phosphorylation or stability of DBC1, we mutated T897 to A897 (Fig. S3 b). We found T897A mutation did not reverse the decrease in protein levels of DBC1 (Fig. S3 c) and thus the site was not significant in the context we had supposed. Nonetheless, the results showed that LKB1 was significantly down-regulating the expression of DBC1. The role of LKB1 in down-regulating DBC1 protein was further confirmed by using LKB1-kinase dead (LKB1-KD) mutant (Fig. 3e). Similar results were obtained with stable cell lines overexpressing Flag tagged LKB1 (pBABE-puro-Flag-LKB1) in U2OS and HeLa cell lines (Fig. 3f). Expression of LKB1 in these cells was confirmed by RT-qPCR (Fig. S4 a,b). We particularly chose HeLa cells as these are known to be LKB1 null cells (Hela -/-) and hence must show higher DBC1 levels in comparison to U2OS cells. Consistent with these data, stable overexpression of LKB1 substantially decreased DBC1 levels (Fig. 3f). In light of above results that LKB1 but not AMPK degrades DBC1, we stimulated cells with various AMPK agonists. Stimulation with AICAR, 2-deoxyglucose (2-DG) and metformin did not affect the protein levels of DBC1 (Fig. S3 d). In summary, using both transiently transfected and stable cell lines, we identified LKB1 as a regulator of DBC1. However, we failed to observe a direct interaction between LKB1 and DBC1 in LC-MS-MS (Fig. 1b) and immunoprecipitation results (data not shown).

**Fig. 3:**
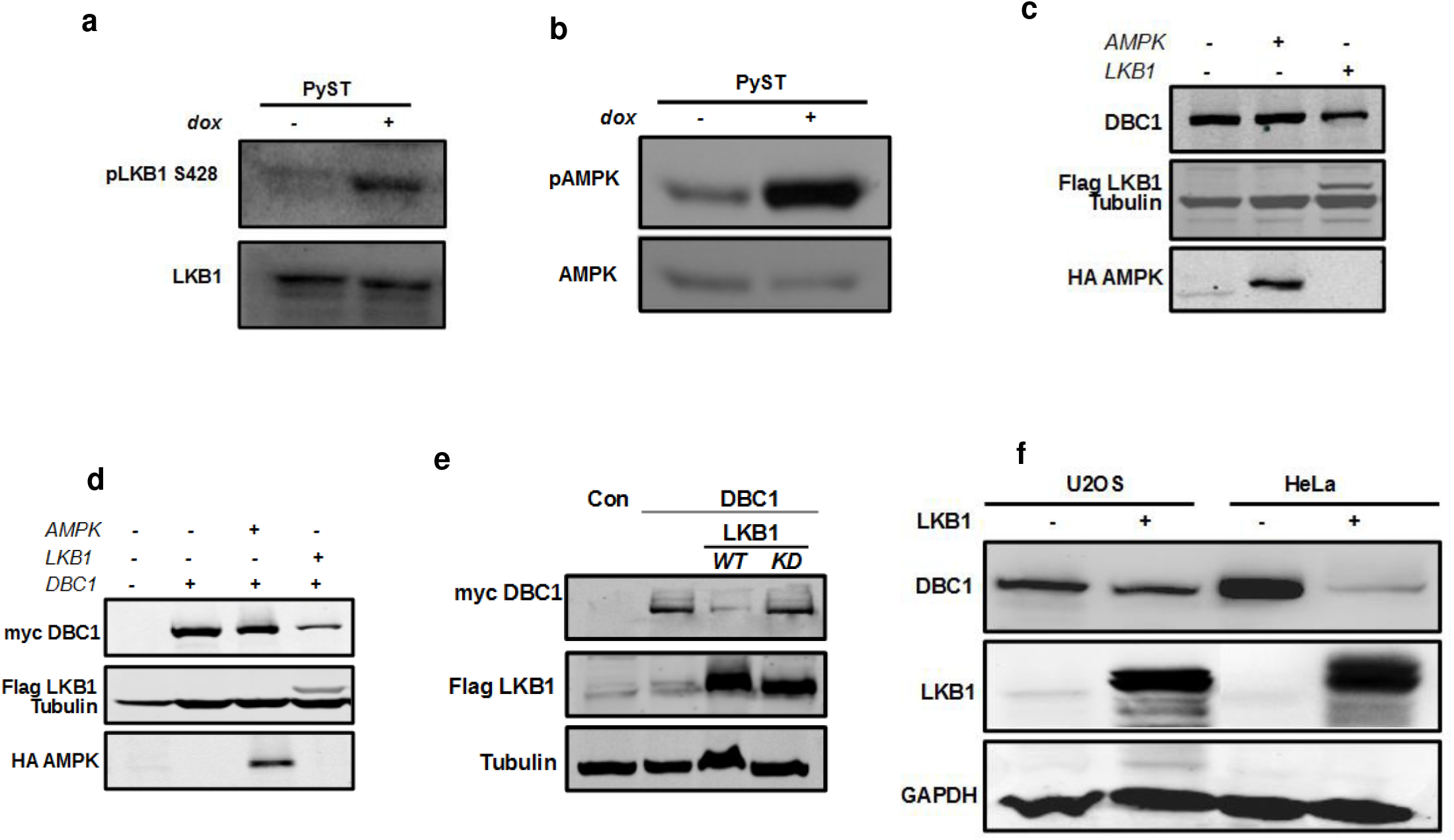
LKB1 is a negative regulator of DBC1. (a, b) PyST activates LKB1: pTREX-PyST cells were induced for 24 hours. Cell lysates were blotted for endogenous pLKB1 and pAMPK protein levels using respective antibodies. (c, d) LKB1 but not APMK caused a significant down-regulation of DBC1 protein amounts at endogenous and exogenous levels respectively. 293T cells were transfected with myc-DBC1, HA-AMPK and Flag-LKB1 constructs. Cells lysates were obtained after about 48 hrs and analyzed by Western blotting using indicated antibodies. (e) 293T cell lines were transfected with myc-DBC1, Flag-LKB1 (WT) and kinase dead (KD) mutant of Flag-LKB1. Western blot of lysates was done using anti-myc for DBC1 and anti-Flag antibody for LKB1. (f) Effect of LKB1 over-expression on DBC1 in stable cell lines. Lysates from U2OS and HeLa cells from control and overexpressing LKB1 were blotted for endogenous DBC1 using anti-DBC1 antibody.

### LKB1 negatively regulates AKT1 via DBC1 and TRB3

DBC1 is known to act as a positive regulator of AKT1 S473 phosphorylation via Tribbles3 (TRB3) (11). It does so by transcriptional repression of TRB3 expression. In absence of growth signals, TRB3 binds to and sequesters AKT1 and prevents its phosphorylation and activation (12). Since expression of PyST causes down-regulation of DBC1, it meant that PyST expressing cells should have higher amounts of TRB3 mRNA and reduced phosphorylation of AKT1 at S473. We had performed whole genome microarray analysis on mRNAs obtained from the inducible PyST expressing cells (pTREX-POLST-HA-Flag), using Human Genome U133 Plus 2 Array (manuscript communicated separately). Microarray data revealed that PyST expressing cells (+dox) had about 2.8 fold increase in TRB3 mRNA levels (Fig. 4a). Our results using RT-qPCR confirmed that PyST expressing cells (+dox) had 5 fold increase in TRB3 mRNA (Fig. 4b). Western blotting results in PyST expressing cells also showed an inverse relationship between DBC1 and TRB3 protein levels (Fig. 4c). Consistent with these results, in DBC1 overexpressing stable cell lines (Fig. 4d, 4e), mRNA levels of TRB3 were decreased by about 5 fold (Fig. 4f). In further support of our data, we observed that over-expression of LKB1, resulted in about 8 fold increase in the mRNA levels of TRB3 (Fig. 4g).

**Fig. 4:**
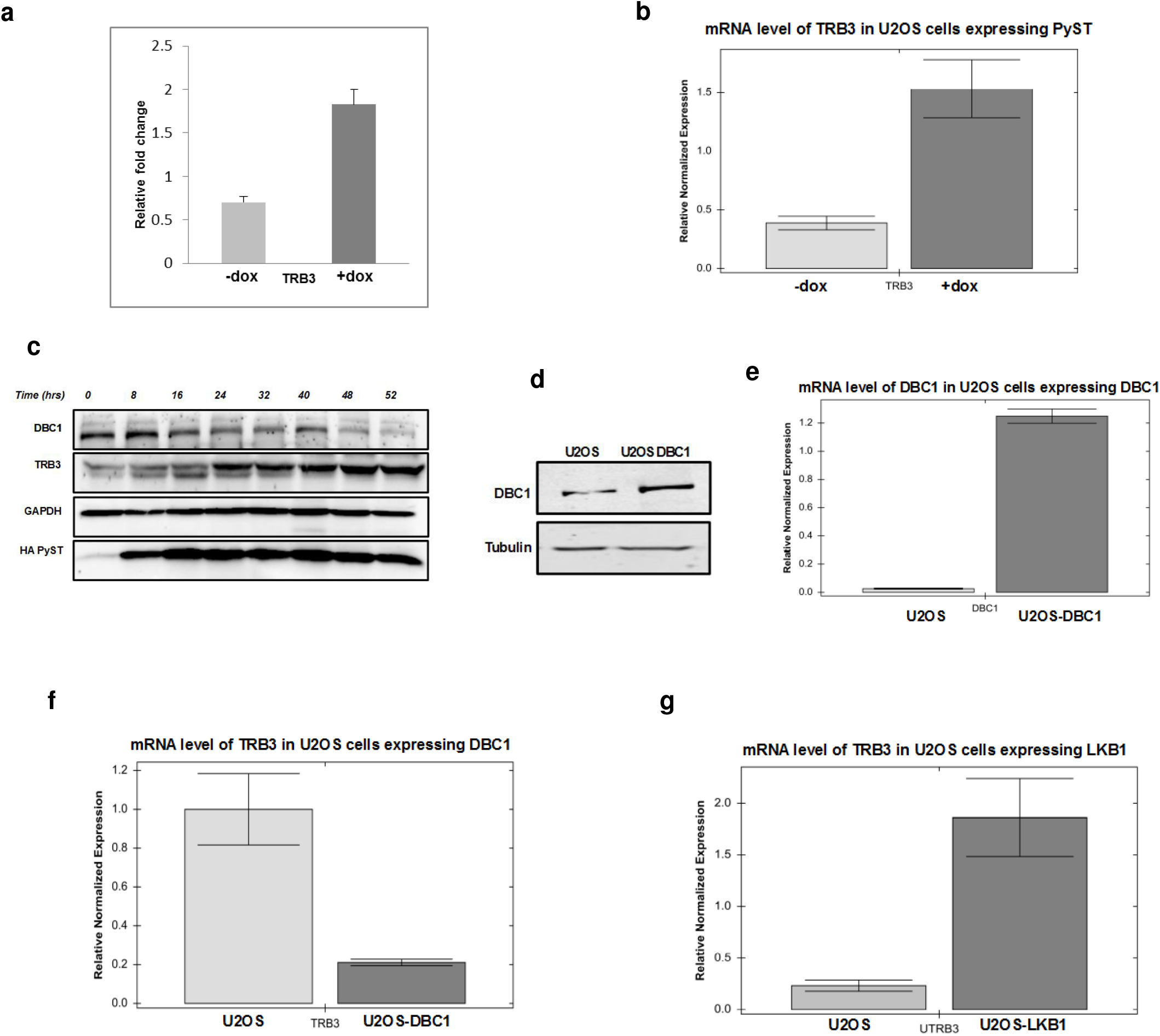
LKB1 positively regulates TRB3 via DBC1. (a) Graphical representation of microarray data from control (-dox) and PyST expressing U2OS cells (+dox) for Tribbles 3. The experiments were carried out as biological triplicates. (b) Fold change for TRB3 mRNA in PyST expressing cells (+Dox, 24 hours) as compared to control cells (-Dox) as shown by RT-qPCR. (c) PyST increases protein levels of TRB3. pTREX-PyST cells were induced for expression of PyST and cell lysates were then analyzed by Western blotting for DBC1, TRB3 and PyST using antibodies against respective proteins. (d, e) DBC1 was stably expressed in U2OS cells and expression at protein and mRNA levels was checked by Western blotting using anti DBC1 antibody and RT-qPCR respectively. (f) Effects of overexpression of DBC1 on TRB3 mRNA levels using RT-qPCR. (g Fold change in the levels of TRB3 mRNA in LKB1 overexpressing cells. For all these experiments, total cellular RNA was extracted from respective cell lines. Following cDNA synthesis, RT-qPCR was carried out using appropriate primers and normalization was done using actin as control.

In light of these results, we wanted to see the impact of PyST expression on AKT1 phosphorylation in pTREX-PyST HA Flag cells. Western blotting showed that expression of PyST led to decreased AKT1 phosphorylation at S473 and T308 (Fig. 5a). We have previously reported that PyST expression causes decreased AKT1 phosphorylation at S473 residue but the mechanism was not studied (7). If LKB1, DBC1 and TRB3 were truly connected, LKB1 and DBC1 would have opposite impacts on AKT1 phosphorylation. It was observed that phosphorylation of AKT1 on S473 was substantially higher in DBC1 over-expressing cell lines (Fig. 5b). Expectedly, the results obtained for LKB1 over-expressing U2OS cells were opposite to those obtained with DBC1 cell lines. LKB1 over-expression, as was already seen (in Fig. 3e), caused down-regulation of DBC1 and a concomitant decrease in AKT1 S473 phosphorylation in both U2OS and HeLa stable cell lines expressing LKB1 (Fig. 5c). To test directly whether LKB1 affects the phosphorylation status of AKT1 as in stable cell lines, we co-expressed AKT1 and LKB1 in 293T cells. Overexpression of LKB1 considerably decreased phosphorylation levels of AKT1 at S473 and T308 residues, thus further confirming the inhibition of AKT1 by LKB1 (Fig. 5d). To confirm that LKB1 truly inhibited AKT1 activity as well, we checked effect of over-expression of LKB1 on forkhead protein (FOXO3a) in 293T cells, using AKT1 and DBC1 as positive controls. FOXO proteins are transcription factors that are well known to be the direct substrates of AKT1. Results confirmed that over-expression of LKB1 caused significant decrease in pFOXO3a signals (Fig. 5e), in response to inhibition of AKT1 by LKB1.

**Fig. 5:**
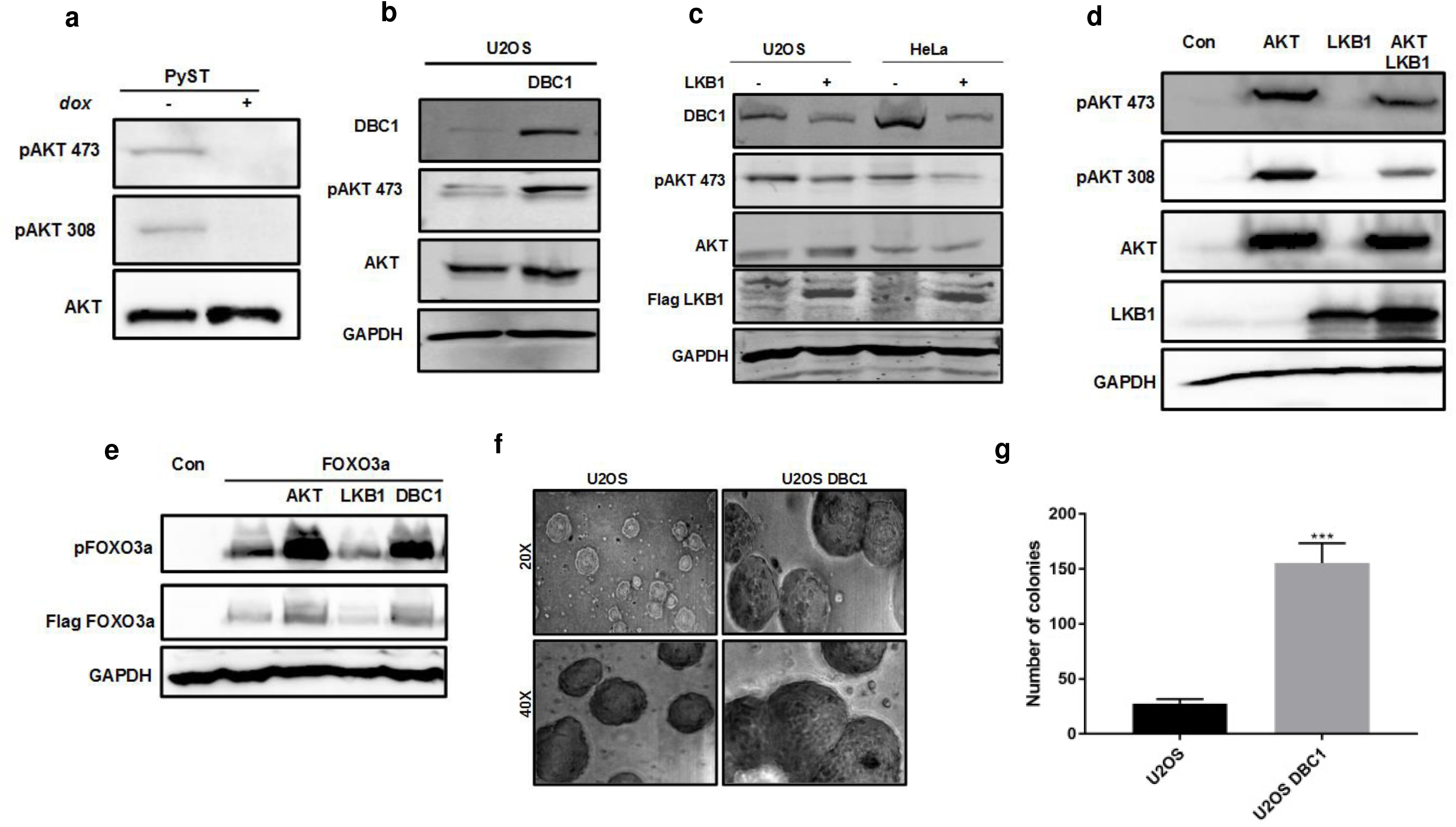
LKB1 regulates AKT1 via DBC1 and TRB3. (a) PyST inhibits phosphorylation of AKT1. Lysates from U2OS cells stably expressing PyST and DBC1 were blotted with antibody specific for phosphorylation of S473 and T308 residues on AKT1. Membrane was stripped and blotted for total AKT protein levels. (b) Effect of DBC1 overexpression on AKT1 phosphorylation. Protein lysates from U2OS cells stably expressing DBC1 were blotted for AKT1 S473 phosphorylation. (c) Effect of LKB1 overexpression on AKT S473 phosphorylation in U2OS and HeLa (LKB1-/-) cell lysates. (d) 293T cells were transiently transfected with AKT1 and LKB1 constructs. pAKT 473, pAKT 308, total AKT and LKB1 were detected in Western blotting using respective antibodies. (e) 293T cells were transiently transfected with pcDNA3-FLAG-FOXO3a and AKT1, LKB1 or DBC1. Cell lysates were then blotted for pFOXO3a and total FOXO3a using indicated antibodies. (f) Soft agar assay for U2OS cells stably over-expressing DBC1 was done in triplicates. Pictures of colonies as seen in figure were taken after 3 weeks of growth. (g) Quantitative representation of soft-agar assay. After 4 weeks of growth, colonies were stained with iodonitrotetrazolium for overnight. Following staining, colonies of 2mm size or bigger were counted and data was analyzed by GraphPad Prism 7 (*P<0.05, **P<0.01, ***P<0.001).

### DBC1 enhances cellular transformation via positive regulation of AKT1

Based on our results especially with regard to activation of AKT1, a well-known proto-oncogene, DBC1 is expected to be a positive regulator of cellular proliferation. We therefore wanted to explore the cellular role of DBC1 independent of its interaction with PyST, with major focus on mitosis and transformation. We checked the ability of DBC1 over-expressing cells to form colonies in soft agar, a gold standard assay for testing the transforming property of the cells. U2OS being a cancerous cell line was expected to form colonies in soft agar. Our results showed that U2OS cells over-expressing DBC1 formed larger and more number of colonies as compared to U2OS cells by themselves (Fig. 5f, g). Since there is a good correlation between *in vitro* transformation and *in vivo* carcinogenesis, these results implicate that DBC1 promotes cellular transformation.

### LKB1 is one of the important requirements for PyST induced mitotic catastrophe

To further validate the importance of LKB1 activation in PyST induced apoptosis, we stably expressed PyST in HeLa cell line (LKB1-/-) and confirmed its expression (Fig. S4c). Western blots showed that PyST expression enhanced phosphorylation of LKB1 at S428 and led to the down-regulation of DBC1 in U2OS but not in HeLa (LKB1 -/-) cells (Fig. 6a), which are known to have genetic ablation of LKB1 alleles. While doxycycline addition promoted mitotic arrest and apoptosis in pTREX-PyST U2OS cells, pTREX-PyST-HeLa cells did not show any such phenotype under similar conditions (Fig. 6b). These results were further confirmed by crystal violet staining of these cells (Fig. 6c) as well as by FACS analysis (Fig. 6d and 6e). In summary, all these results support our argument that LKB1 plays an important role in mediating the effect of PyST in causing mitotic arrest and apoptosis.

**Fig. 6:**
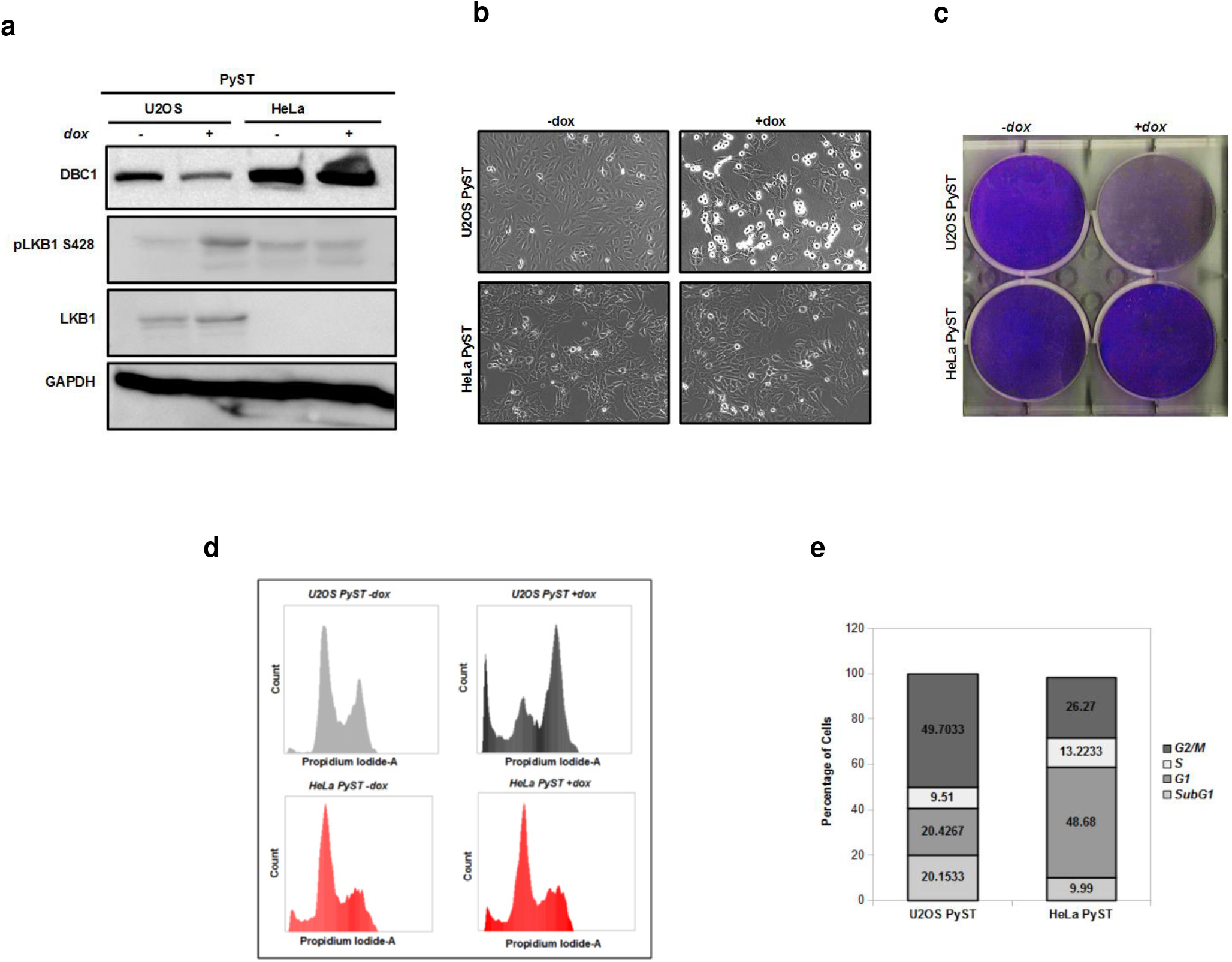
LKB1 is important for PyST mediated mitotic arrest and cell death. (a) Effect of PyST over-expression on DBC1 protein levels. U2OS and HeLa cells over-expressing PyST were blotted for endogenous DBC1 using anti-DBC1 antibody. (b) Stable over-expression of PyST in U2OS and HeLa cells. Cell lines were induced for PyST expression and phenotype was observed and imaged using a simple microscope. (c) 48 hours post-induction, cells were stained with crystal violet stain and plates were imaged. (d) Flow cytometric analysis of U2OS and HeLa cells over-expressing PyST. Cell cycle analysis was done by using BD FACSverse. (e) Data from FACS for +dox samples as expressed in graphical form.

**Figure 7:**
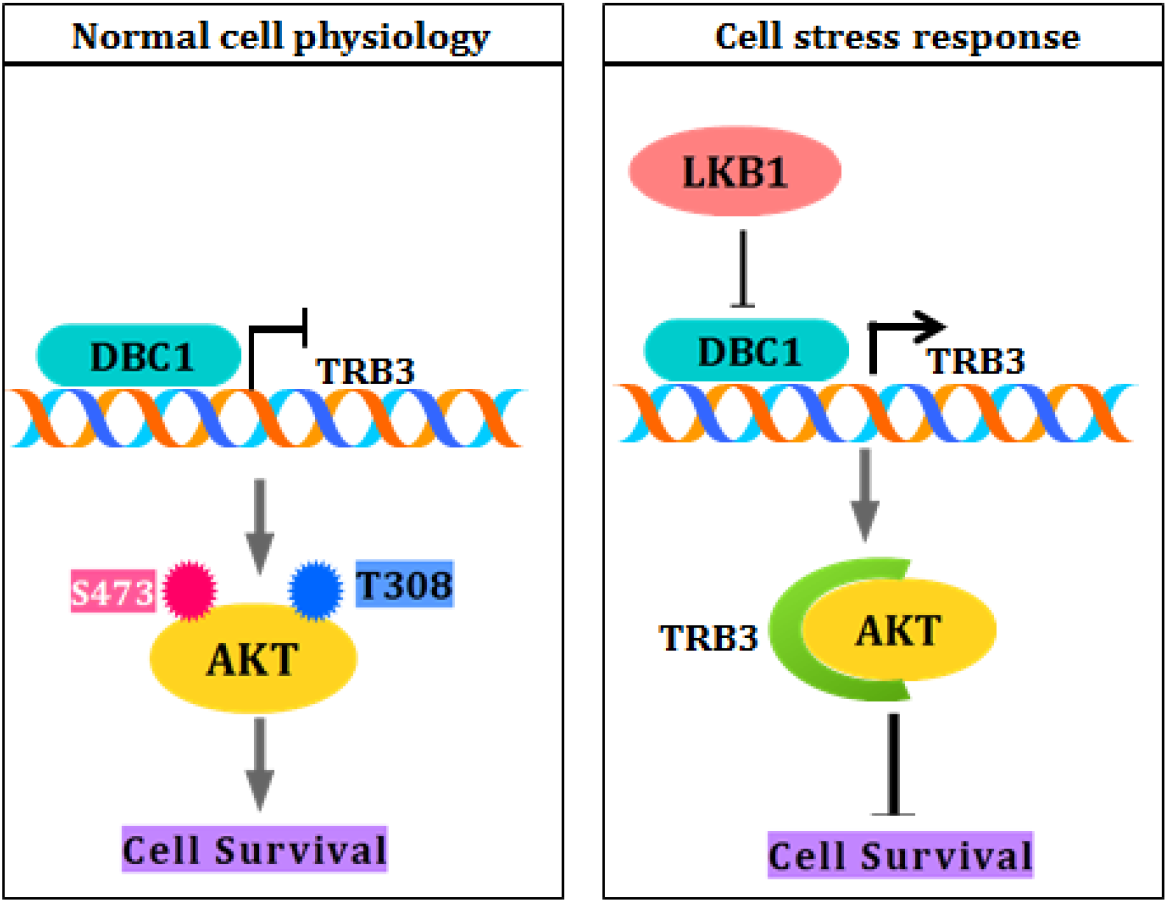
Model showing connection between LKB1, DBC1, TRB3 and AKT1. Under normal cellular conditions, DBC1 transcriptionally represses TRB3. In absence of TRB3, AKT is maintained in phosphorylated and activated form leading to cell survival. In response to energy deficit, LKB1 is activated. Once active, LKB1 leads to down-regulation of DBC1. Absence of DBC1 releases transcriptional inhibition of TRB3. TRB3 sequesters AKT1 and prevents its phosphorylation and activation. Once AKT1 is inhibited, major pro-survival cellular signals are shut, thereby restricting cell survival.

### Energy deficit is critical for the induction of mitotic arrest and apoptosis by PyST

PyST expression causes extensive poly ADP-ribosylation (PARylation) of various cellular proteins in host cells (7), thus causing severe energy deficit in these cells. If energy stress is important for PyST phenotype, then supplementing these cells with exogenous NAD^+^ should alleviate the apoptotic phenotype of PyST expressing cells. To test that, we treated PyST expressing cells with 250 µM of NAD^+^. FACS analysis of these cells showed that NAD^+^ addition rescued these cells from cell death and prevented mitotic arrest (Fig. S4d, e). This again supported our observation that activation of energy stress pathway and the consequent stimulation of LKB1 activity play a pivotal role in mediating PyST induced cell death.

Our results therefore indicate a logical link between LKB1, DBC1, TRB3 and AKT1. Consequent to cellular stress (PyST expression), activation of LKB1 leads to decreased DBC1 protein amounts, which enhances TRB3 mRNA expression. Increased TRB3 amounts leads to the sequestering of AKT1, thus preventing its phosphorylation and activation by upstream kinases (Graphical Abstract). This results in the inhibition of AKT1 mediated pro-survival signaling pathway.

## Discussion

In this study, using polyoma small T antigen expressing cell lines, we delineate a molecular mechanism showing that LKB1 negatively regulates AKT1. LKB1 acts as a tumor suppressor in cells and is known to negatively regulate PI3 kinase pathway through activation of AMP kinase (13). Both LKB1 and AMP kinase are sensors of cellular energy metabolism and are activated in response to low cellular energy levels (14). In contrast, AKT1 is a well-known oncogene that operates downstream of the PI3 kinase pathway and is activated in response to abundant nutrients and growth factors, thus promoting cell growth, division and survival (15). Both AKT1 and LKB1 genes are often mutated in cancers, leading to tumorigenesis.

Our results suggest that expression of small T causes severe energy deficit in the host mammalian cells, which activates LKB1 as was seen by enhanced LKB1 phosphorylation. Activated LKB1, through an unknown mechanism, leads to the degradation of DBC1 protein. DBC1 is a transcriptional co-repressor for TRB3 expression (11). In the absence of growth factors, Tribbles3 is known to sequester AKT1 and prevent its phosphorylation and activation by upstream kinases (11, 12). Since activated LKB1 degrades DBC1, this would mean upregulation of TRB3 expression and consequent inactivation of AKT1. Using both indirect (in PyST overexpressing cell lines) and direct approaches (LKB1 and AKT1 co-transfections), our results support a mechanism showing that activated LKB1 negatively regulates AKT1 through DBC1 and TRB3. This is a novel finding showing how LKB1 regulates the activity of AKT1 in response to cellular energy status. This obviously raises the question as to how LKB1 is activated in PyST expressing cells. Expression of PyST causes overwhelming mitotic arrest in the host mammalian cells(8). These cells lack chromosomal congression and proper spindle formation during mitotic progression, which stimulates PARP1 activity, thereby activating DNA repair pathways in these cells. This leads to extensive ADP ribosylation (called PARylation) of numerous cellular proteins. We have previously shown that PyST expressing cells have abnormally high levels of ADP Ribosylation (7). We attributed the effect of LKB1 activation by PyST to occur via inhibition of PP2A, a known phosphatase for LKB1. However, quite surprisingly, using different mutants, we showed that PyST activates LKB1 through its N-terminal J domain, which is known to bind and activate heat shock proteins. What is the exact mechanism for this activation remains an interesting question to be to be explored. We presume that PyST would bring together heat shock proteins, DBC1, LKB1 and proteasomes in close proximity on its J domain, thus provoking LKB1 to degrade DBC1. Support for this assumption comes from the papers that have shown interaction between DBC1 and heat shock proteins (16) as well as between heat shock proteins and proteasomal complex (Luders et al., 2000).

Interestingly, we also identified LKB1 as a negative regulator of DBC1. LKB1 causes DBC1 downregulation and while most of the effects of LKB1 occur via AMPK, our study reports the otherwise. It has been previously reported that LKB1 plays a role in mitotic regulation in hematopoietic cells (HSCs) independently of AMP kinase. These cells from LKB1 -/- mice showed various abnormalities like supernumerary centrosomes, abnormal spindles and aneuploidy (17, 18). Another study showed that LKB1 regulates mitosis in an AMPK independent manner by promoting dephosphorylation of PLK1, an important protein involved in mitotic progression(19). Consistent with these reports, our results have also shown that LKB1 degrades DBC1 in an AMPK independent manner. We believe that LKB1 may either directly phosphorylate DBC1 or indirectly through one of the 13 known downstream kinases other than AMPK.

The role of DBC1 in human cancer has been largely paradoxical. In our study, we found that DBC1 promotes cellular transformation in vitro. Recently it has been reported that DBC1 has tumor suppressor activity by stabilizing p53 (20). However, taken together, our results particularly with increased AKT1 signaling and anchorage independence assay show that DBC1 over-expression in U2OS cells promotes oncogenic pathways. These results are consistent with some recent reports also suggesting the oncogenic role of DBC1 (21–23).

Future studies are required to elucidate the mechanism behind down-regulation of DBC1 by PyST and LKB1. It also remains to be fully elucidated how LKB1 is specifically activated by PyST and whether LKB1 activation is the only pre-requisite for PyST induced mitotic catastrophe. It would be interesting to analyze whether DBC1 overexpression is associated with patient survival and treatment outcomes in cancer patients.

## Acknowledgements

Shaida Andrabi was supported by the Department of Biotechnology (DBT), India through the following grants: Ramalingaswami Fellowship, BT/PR5743/BRB/10/1100/2012, BT/PR13605/MED/30/1525/2015 and BT/PR2283/AGR/36/692/2011, and DST-FIST grants. We also thank DST, India for providing INSPIRE fellowships to Zarka Sarwar, Sameer Bhat, Qaaifah Gillani and also to CSIR-UGC for providing fellowship to Irfana Reshi and Misba Shah. We thank Prof. Shakil Wani, Dr. Zahid and their colleagues for allowing us to carry out the FACS analysis at SKUAST campus, Shuhama, Srinagar. We also want to thank Prof. Thomas Roberts, Dr. Ole Gjoerup and Prof. Brian Schaffhausen for reviewing this manuscript.

## Author details

ZS and SA designed the experiments, interpreted the data and wrote the manuscript. ZS carried out the experiments. SB, QG, IR and MS carried out some of the experiments. GA and JM carried out the LC-MS-MS experiments, compiled the data and helped with interpretation of the MS results.

## Material and Methods

### Materials

All antibodies were purchased from Cell signaling technology, USA (CST). Tribbles 3 antibody was purchased from Sigma. Primers for cloning and RT qPCR were supplied by Integrated DNA Technologies (IDT). Kits for cDNA synthesis and RT qPCR were purchased from BioRad, USA (iScriptcDNA Synthesis Kit and Sso Advanced Universal SYBR Green Supermix). Cell culture reagents like DMEM, FCS and antibiotics were purchased from Invitrogen/Thermofisher Scientific, USA.

### Cloning

myc-tagged DBC1 (myc-DBC1) cloned in pcDNA3 was obtained from Addgene, USA. For making stable cell lines, DBC1 was cloned in retroviral vector pWZL-Blast using EcoR I and Xho I sites, which were compatible between two vectors.

### Transient transfections

For transient transfections, polyethylenimine (PEI) method was routinely used. Briefly, 293T cells were grown at a confluence of 60-70% in 60mm plates. A total of 5 µg of the DNA was mixed with 400 µl of DMEM and then 15 µl of PEI was added drop wise. The mixture was incubated for 15-20 mins and was then added slowly to the plates and incubated overnight. 16 hours post transfection, the plates were washed with phosphate-buffered saline (PBS), and then supplemented with fresh DMEM [Dulbecco’s modified Eagle’s medium (DMEM) containing 10% fetal calf serum and 1% Pencillin-Streptomycin]. 48 hours post transfection, the cells were harvested, and the extracts were used for further experiments.

### Retroviral/lentiviral infections

Phoenix cells stably expressing packaging proteins were transiently transfected with pWZL Blast DBC1 using PEI as a transfecting reagent. Briefly, 3 µg pWZl-DBC1, 1 µg gag-pol and 1 µg of VSVG were mixed in 400 µl of DMEM and 15 µg of PEI was added drop wise. This transfection mixture was then added to cells grown in 60-mm plates at a confluence of 60 to 80%. Next day, cells were washed with (PBS) and fresh DMEM was added. After 48 hours, cells that were split a day before were infected with the viral supernatants obtained from the above transfected cells and supplemented with 8 µg of polybrene/ml (Sigma) for at least 6 hours. This process was repeated the next day also. For selection of DBC1 expressing cells, cells were grown in the presence of blasticidin (10 µg/ml). For making double stable cell lines for PyST and DBC1, U2OS cells stably expressing pTREX-PyST-HA-Flag were infected with viral titer generated by using pWZL-blast-DBC1. Cells were selected using puromycin (5 µg/ml) for PyST and blasticidin for DBC1 (10 µg/ml). Similarly, for making stable LKB1 overexpressing cell lines, viral titer was generated in Pheonix cells using pBABE-puro-Flag-LKB1. U2OS and HeLa cells were infected with the viral supernatant and cells were selected using puromycin (5 µg/ml for U2OS and 2 µg/ml for HeLa). For making HeLa cells stably expressing PyST, FT cells expressing packaging proteins were transiently transfected with pTREX-PyST-HA-Flag using PEI as a transfecting reagent. 2.5 µg pTREX-PyST, 2.25 µg R and 0.25 µg of VSVG were mixed in 400 µl of DMEM and 15 µl of PEI was added drop wise. Viral titer thus generated was used to infect HeLa cells and following infection, the cells were selected with puromycin (2.5 µg/ml).

### Western blotting

Cells were washed with ice cold PBS, harvested and resuspended in lysis buffer [Tris-Cl 50 mM, NaCl 150 mM, Glycerol 10%, SDS 0.1%, NP-40 1%, EDTA 2 mM, supplemented with protease and phosphatase inhibitors (Complete, Roche)]. Equal amounts of protein were loaded, as estimated by Bradford assay (Bio-Rad Laboratories).

### Real time quantitative PCR (RT-qPCR)

For RT-qPCR, cells were grown in 60 mm dishes. RNA was isolated by using Trizol method. cDNA synthesis was carried out using Bio-Rad iScript cDNA Synthesis Kit. Bio-Rad Sso Advanced Universal SYBR Green Supermix was used for RT-qPCR and data was analysed by Bio-Rad CFX Manager. Following primer pairs were used for RT-qPCR: DBC1 5’-AAGGTGCAAACGCTCTCCAACCAG-3’ and 3’-GGATGTTTGGAAGAGACTCAGAG-5’, Actin 5’-TTAGTTGCGTTACACCCTTTC-3’ and 3’-ACC TTC ACC GTT CCA GTT T-5’, TRB3 5’-TGCCCTACAGGCACTGAGTA-3’ and 3’-GTCCGAGTGAAAAAGGCGTA-5’, LKB1 5’-ACTGAGGAGGTTACGGCACA-3’ and 3’-CCTGGCACACTGGGAAAC-3’.

### Cell cycle analysis by fluorescence-activated cell sorting (FACS)

U2OS cells stably expressing DBC1 and polyoma small T were grown to a confluence of 60-70% in 60 mm plates. The cells were washed with PBS/EDTA (0.1%) and then incubated at 37°C for 5 minutes in PBS/EDTA, scrapped from the plate and collected in 15 ml falcon tubes. The cells were pelleted at 1000 xg for 5 minutes and washed once with PBS/serum (1%). Finally, the cells were fixed by adding 10 ml of cold ethanol while vortexing to prevent clumping. The fixed cells were stored at 4°C at least overnight in the dark. Prior to acquisition on Flow Cytometer, the cells were centrifuged at 1,000xg for 5min and then washed once with PBS/serum (1%). Pellets were resuspended in 500 µl of propidium iodide-RNase solution (50 µg/ml propidium iodide, 10mM Tris, pH 7.5, 5 mM MgCl_2_, and 20µg/ml RNase A), and incubated at 37°C for 30 min. Samples were then acquired by using BD FACSverse and analyzed by BD FACSverse software. Statistical significance was calculated by using GraphPad Prism 7.0 software.

### Soft-agar assay

Soft-agar assays were done in 60 mm dishes in triplicates. A stock of 1.8% agar (Noble agar; Sigma) was autoclaved and stored at room temperature. For bottom agar, 0.6% agar in DMEM supplemented with 0.1% FBS was used. For top agar, briefly, 1×10 were suspended in minimal essential growth medium (DMEM with 0.1% FBS and 10% penicillin/streptomycin) containing 0.33% agar and incubated for 3 weeks. The plates were fed with 500µl of DMEM twice a week. Plates were analyzed for colony number and size. Size and Statistical analysis were done using GraphPad Prism 7.0.

### Immunoprecipitation

293T cells were grown in 60 mm plates, washed once with cold PBS, harvested, and lysed in 100 µl of lysis buffer in the presence of protease and phosphatase inhibitors for 30 min on ice. The extracts were spun down at 10,000 rpm for 10 min. 20 µl of IgG sepharose beads were premixed with 5 µl of antibodies for 30 min to 1 h. About 20 µl of the lysate was aliquoted for the whole cell lysate blot (WCL). Equal amounts of protein as determined by Bradford assay were then added to the beads-antibody mixture and were kept on rotator overnight at 4°C. The mixtures were then washed twice with cold PBS followed by a final wash with cold water. The beads were mixed with 2X sample buffer, boiled, and the proteins were analyzed by Western blotting.

### Site directed mutagenesis

Threonine to Alanine mutation at 897 position of DBC1 (T897A) was carried out using NEB Q5 High Fidelity DNA polymerase and protocol for the same was followed as per the instructions of the manufacturer (New England Biolabs). Following PCR, Dpn I treatment was carried and the PCR mixture was transformed into DH5α cells. DNA from the colonies was isolated using miniprep kit and mutants were confirmed by DNA sequencing.

### Crystal violet staining

For growth assays by crystal violet staining, cells were grown in 60mm dishes. The assays were carried out in duplicates. At desired time points, the cells were washed with PBS, fixed with methanol for 10 mins and then stained with 0.1% crystal violet for 15 minutes. Excess stain was removed by washing with water. For quantification purposes, the cells were then destained by adding 2 ml of 10% acetic acid to each well and incubated for 20 minutes on a shaker. The destained solutions were collected and absorbance was measured at 590 nm.

### Microarray

Total cellular RNA was isolated from pTREX-PyST cells (in triplicates) at about 20 hours of doxycycline addition using Trizol method (Invitrogen). RNA was purified using RNAeasy kit (Qiagen) as per manufacturers specifications used for microarray analysis, using Affymetrix Human Genome U133 Plus 2 Array chips. Microarray experiments were performed at DFCI, Harvard Medical School microarray core facility. Gene expression changes with >2-fold alterations were considered significant.

### Mass spectrometry analysis of TAP samples: Mass spectrometry analysis of TAP samples

Purified protein complexes were analyzed by mass spectrometry (LC/MS-MS) as described (24) with minor modifications:

Proteins from the TAP samples were directly processed in solution: Cysteine residues were reduced with 10 mM DTT for 30 minutes at 56°C in presence of 0.1% RapiGest SF (Waters). Proteins were digested overnight at 37ºC using 5 µg of trypsin after adjusting the pH to 8.0 with Tris buffer. Cys-HA peptide was removed from the TAP samples by incubating the digests with 20 µL of a 50% slurry of Activated Thiol Sepharose 4B (GE healthcare) for 30 minutes at room temperature. The HA peptide-depleted solutions were acidified by adding TFA to a final concentration of 1%, desalted by batch-mode reverse phase (RP, Poros 10R2, Applied Biosystems) solid phase extraction and concentrated in a vacuum concentrator. Peptides were solubilized in 0.1% Formic Acid containing 25% Acetonitrile and further purified by Strong Cation Exchange (SCX Poros 10HS, Applied Biosystems). Peptides were sequentially eluted with 0.1% Formic Acid containing 25% Acetonitrile and the following concentrations of KCl: 10 mM, 30 mM, 50 mM, 100 mM, 150 mM, 250 mM and 300 mM. Fractionated peptides were concentrated in a vacuum concentrator, and reconstituted with 20 µl of 0.1% TFA. Peptides from each fraction were loaded onto a precolumn (4 cm POROS 10R2, Applied Biosystems) and eluted with an HPLC gradient (NanoAcquity UPLC system, Waters; 2%–35% B in 45 min; A = 0.2 M acetic acid in water, B = 0.2 M acetic acid in acetonitrile). Peptides were resolved on a self-packed analytical column (12 cm Monitor C18, Column Engineering) and introduced in the mass spectrometer (LTQ-Orbitrap-XL mass spectrometer, Thermo) equipped with a Digital PicoView electrospray source platform (25) (New Objective). The spectrometer was operated in data dependent mode where the top 8 most abundant ions in each MS scan were subjected to CAD (35% normalized collision energy) and subjected to MS2 scans (isolation width = 2.8 Da, threshold = 20,000). Dynamic exclusion was enabled with a repeat count of 1 and exclusion duration of 30 seconds. ESI voltage was set to 2.2 kV.

MS spectra were recalibrated using the background ion (Si(CH3)2O)6 at m/z 445.12 +/-0.03 and converted into a Mascot generic file format (.mgf) using multiplierz scripts (25, 26). Spectra were searched using Mascot (version 2.6) against two databases consisting of human protein sequences (downloaded from Uniprot on 09/27/2017) and common lab contaminants, with automatic Mascot Decoy search enabled. Precursor tolerance was set to 15ppm and product ion tolerance to 0.6 Da. Search parameters included trypsin specificity, up to 2 missed cleavages, fixed carbamido-methylation (C, +57 Da) and variable oxidation (M, +16 Da). Spectra matching to peptides from the reverse database were used to calculate a global false discovery rate, and were discarded. Data were further processed to remove peptide spectral matches (PSMs) to the forward database with an FDR greater than 1.0%. A fast peptide matching algorithm was used to map peptide sequences to all possible human genes in the search database (26). Peptides shared by two or more genes were excluded from consideration when constructing the final protein list. Any protein identified in more than 1% of 108 negative TAP controls (27) were removed from the sets of interactors. Proteins identified in the control TAP experiments were also removed. where the top 8 most abundant ions in each MS scan were subjected to CAD (35% normalized collision energy) and subjected to MS2 scans (isolation width = 2.8 Da, threshold = 20,000). Dynamic exclusion was enabled with a repeat count of 1 and exclusion duration of 30 seconds. ESI voltage was set to 2.2 kV.

MS spectra were recalibrated using the background ion (Si(CH3)2O)6 at m/z 445.12 +/-0.03 and converted into a Mascot generic file format (.mgf) using multiplierz scripts. Spectra were searched using Mascot (version 2.6) against two databases consisting of human protein sequences (downloaded from Uniprot on 09/27/2017) and common lab contaminants, with automatic Mascot Decoy search enabled. Precursor tolerance was set to 15ppm and product ion tolerance to 0.6 Da. Search parameters included trypsin specificity, up to 2 missed cleavages, fixed carbamido-methylation (C, +57 Da) and variable oxidation (M, +16 Da). Spectra matching to peptides from the reverse database were used to calculate a global false discovery rate, and were discarded. Data were further processed to remove peptide spectral matches (PSMs) to the forward database with an FDR greater than 1.0%. A fast peptide matching algorithm was used to map peptide sequences to all possible human genes in the search database. Peptides shared by two or more genes were excluded from consideration when constructing the final protein list. Any protein identified in more than 1% of 108 negative TAP controls were removed from the sets of interactors. Proteins identified in the control TAP experiments were also removed.

### Immunofluoresence

U2OS and HeLa cells were grown on coverslips upto 50% confluence. Cells were washed with chilled PBS and fixed with 4% paraformaldehyde (PFA) in PBS. Cells were permeabilized with 0.15 v/v% Triton-X 100 for 3 minutes. Nonspecific sites were blocked with 2% bovine serum albumin (Sigma-Aldrich) in PBS for 45 mins. Fixed cells were stained with the following antibodies (for 3 hrs at 37°C): β-Tubulin (1:100, Sigma-Aldrich), DBC1 (1:100, Cell signaling), HA (1:100, Invitrogen). Primary antibodies were visualized by secondary antibodies conjugated to Alexa fluor 488 and Alexa fluor 594 (1:5000, Jackson ImmunoResearch) at 37°C for 1 h. Cells were stained with DAPI (BioRad) for 10 minutes. Pictures were taken with an inverted fluorescence microscope (Axio Observer, Colibri7, Carl Zeiss MicroImaging, Inc.) using filters for DAPI, GFP and mRF12, X20, X63, DIC Apotome.2 oil objectives. Pictures were generated with ZEN 2.3 pro (Zeiss).

## References

1. Hamaguchi, M., Meth, J. L., von Klitzing, C., Wei, W., Esposito, D., Rodgers, L., Walsh, T., Welcsh, P., King, M.-C., and Wigler, M. H. (2002) DBC2, a candidate for a tumor suppressor gene involved in breast cancer. Proc. Natl. Acad. Sci. U. S. A. 99, 13647–13652

2. Radvanyi, L., Singh-Sandhu, D., Gallichan, S., Lovitt, C., Pedyczak, A., Mallo, G., Gish, K., Kwok, K., Hanna, W., Zubovits, J., Armes, J., Venter, D., Hakimi, J., Shortreed, J., Donovan, M., Parrington, M., Dunn, P., Oomen, R., Tartaglia, J., and Berinstein, N. L. (2005) The gene associated with trichorhinophalangeal syndrome in humans is overexpressed in breast cancer. Proc. Natl. Acad. Sci. U. S. A. 102, 11005–11010

3. Kim, J.-E., Chen, J., and Lou, Z. (2009) p30 DBC is a potential regulator of tumorigenesis. Cell Cycle Georget. Tex. 8, 2932–2935

4. Hardie, D. G., and Alessi, D. R. (2013) LKB1 and AMPK and the cancer-metabolism link - ten years after. BMC Biol. 11, 36

5. Hemminki, A., Markie, D., Tomlinson, I., Avizienyte, E., Roth, S., Loukola, A., Bignell, G., Warren, W., Aminoff, M., Hoglund, P., Jarvinen, H., Kristo, P., Pelin, K., Ridanpaa, M., Salovaara, R., Toro, T., Bodmer, W., Olschwang, S., Olsen, A. S., Stratton, M. R., de la Chapelle, A., and Aaltonen, L. A. (1998) A serine/threonine kinase gene defective in Peutz-Jeghers syndrome. Nature. 391, 184–187

6. Entius, M. M., Keller, J. J., Westerman, A. M., van Rees, B. P., van Velthuysen, M. L., de Goeij, A. F., Wilson, J. H., Giardiello, F. M., and Offerhaus, G. J. (2001) Molecular genetic alterations in hamartomatous polyps and carcinomas of patients with Peutz-Jeghers syndrome. J. Clin. Pathol. 54, 126–131

7. Andrabi, S., Gjoerup, O. V., Kean, J. A., Roberts, T. M., and Schaffhausen, B. (2007) Protein phosphatase 2A regulates life and death decisions via Akt in a context-dependent manner. Proc. Natl. Acad. Sci. U. S. A. 104, 19011–19016

8. Pores Fernando, A. T., Andrabi, S., Cizmecioglu, O., Zhu, C., Livingston, D. M., Higgins, J. M. G., Schaffhausen, B. S., and Roberts, T. M. (2015) Polyoma small T antigen triggers cell death via mitotic catastrophe. Oncogene. 34, 2483–2492

9. Hwang, J. H., Pores Fernando, A. T., Faure, N., Andrabi, S., Adelmant, G., Hahn, W. C., Marto, J. A., Schaffhausen, B. S., and Roberts, T. M. (2014) Polyomavirus small T antigen interacts with yes-associated protein to regulate cell survival and differentiation. J. Virol. 88, 12055–12064

10. Tian, Y., Li, D., Dahl, J., You, J., and Benjamin, T. (2004) Identification of TAZ as a binding partner of the polyomavirus T antigens. J. Virol. 78, 12657–12664

11. Restelli, M., Magni, M., Ruscica, V., Pinciroli, P., De Cecco, L., Buscemi, G., Delia, D., and Zannini, L. (2016) A novel crosstalk between CCAR2 and AKT pathway in the regulation of cancer cell proliferation. Cell Death Dis. 7, e2453

12. Du, K., Herzig, S., Kulkarni, R. N., and Montminy, M. (2003) TRB3: a tribbles homolog that inhibits Akt/PKB activation by insulin in liver. Science. 300, 1574–1577

13. Shaw, R. J., Bardeesy, N., Manning, B. D., Lopez, L., Kosmatka, M., DePinho, R. A., and Cantley, L. C. (2004) The LKB1 tumor suppressor negatively regulates mTOR signaling. Cancer Cell. 6, 91–99

14. Hardie, D. G., Ross, F. A., and Hawley, S. A. (2012) AMPK: a nutrient and energy sensor that maintains energy homeostasis. Nat. Rev. Mol. Cell Biol. 13, 251–262

15. Vivanco, I., and Sawyers, C. L. (2002) The phosphatidylinositol 3-Kinase AKT pathway in human cancer. Nat. Rev. Cancer. 2, 489–501

16. Kim, W., Cheon, M. G., and Kim, J.-E. (2017) Mitochondrial CCAR2/DBC1 is required for cell survival against rotenone-induced mitochondrial stress. Biochem. Biophys. Res. Commun. 485, 782–789

17. Gan, B., Hu, J., Jiang, S., Liu, Y., Sahin, E., Zhuang, L., Fletcher-Sananikone, E., Colla, S., Wang, Y. A., Chin, L., and Depinho, R. A. (2010) Lkb1 regulates quiescence and metabolic homeostasis of haematopoietic stem cells. Nature. 468, 701–704

18. Gurumurthy, S., Xie, S. Z., Alagesan, B., Kim, J., Yusuf, R. Z., Saez, B., Tzatsos, A., Ozsolak, F., Milos, P., Ferrari, F., Park, P. J., Shirihai, O. S., Scadden, D. T., and Bardeesy, N. (2010) The Lkb1 metabolic sensor maintains haematopoietic stem cell survival. Nature. 468, 659–663

19. Werle, K., Chen, J., Xu, H.-G., Zhao, R.-X., He, Q., Lu, C., Cui, R., Liang, J., Li, Y.-L., and Xu, Z.-X. (2014) Liver kinase B1 regulates the centrosome via PLK1. Cell Death Dis. 5, e1157

20. Qin, B., Minter-Dykhouse, K., Yu, J., Zhang, J., Liu, T., Zhang, H., Lee, S., Kim, J., Wang, L., and Lou, Z. (2015) DBC1 functions as a tumor suppressor by regulating p53 stability. Cell Rep. 10, 1324–1334

21. Kim, H. J., Moon, S. J., Kim, S.-H., Heo, K., and Kim, J. H. (2018) DBC1 regulates Wnt/beta-catenin-mediated expression of MACC1, a key regulator of cancer progression, in colon cancer. Cell Death Dis. 9, 831

22. Moon, S. J., Jeong, B. C., Kim, H. J., Lim, J. E., Kwon, G. Y., and Kim, J. H. (2018) DBC1 promotes castration-resistant prostate cancer by positively regulating DNA binding and stability of AR-V7. Oncogene. 37, 1326–1339

23. Yu, X., Wang, M., Han, Q., Zhang, X., Mao, X., Wang, X., Li, X., Ma, W., and Jin, F. (2018) ZNF326 promotes a malignant phenotype of breast cancer by interacting with DBC1. Mol. Carcinog. 57, 1803–1815

24. Adelmant, G., Calkins, A. S., Garg, B. K., Card, J. D., Askenazi, M., Miron, A., Sobhian, B., Zhang, Y., Nakatani, Y., Silver, P. A., Iglehart, J. D., Marto, J. A., and Lazaro, J.-B. (2012) DNA ends alter the molecular composition and localization of Ku multicomponent complexes. Mol. Cell. Proteomics MCP. 11, 411–421

25. Ficarro, S. B., Zhang, Y., Lu, Y., Moghimi, A. R., Askenazi, M., Hyatt, E., Smith, E. D., Boyer, L., Schlaeger, T. M., Luckey, C. J., and Marto, J. A. (2009) Improved electrospray ionization efficiency compensates for diminished chromatographic resolution and enables proteomics analysis of tyrosine signaling in embryonic stem cells. Anal. Chem. 81, 3440–3447

26. Askenazi, M., Marto, J. A., and Linial, M. (2010) The complete peptide dictionary--a meta-proteomics resource. Proteomics. 10, 4306–4310

27. Rozenblatt-Rosen, O., Deo, R. C., Padi, M., Adelmant, G., Calderwood, M. A., Rolland, T., Grace, M., Dricot, A., Askenazi, M., Tavares, M., Pevzner, S. J., Abderazzaq, F., Byrdsong, D., Carvunis, A.-R., Chen, A. A., Cheng, J., Correll, M., Duarte, M., Fan, C., Feltkamp, M. C., Ficarro, S. B., Franchi, R., Garg, B. K., Gulbahce, N., Hao, T., Holthaus, A. M., James, R., Korkhin, A., Litovchick, L., Mar, J. C., Pak, T. R., Rabello, S., Rubio, R., Shen, Y., Singh, S., Spangle, J. M., Tasan, M., Wanamaker, S., Webber, J. T., Roecklein-Canfield, J., Johannsen, E., Barabasi, A.-L., Beroukhim, R., Kieff, E., Cusick, M. E., Hill, D. E., Munger, K., Marto, J. A., Quackenbush, J., Roth, F. P., DeCaprio, J. A., and Vidal, M. (2012) Interpreting cancer genomes using systematic host network perturbations by tumour virus proteins. Nature. 487, 491–495

